# A potential function for the helicase Dbp5 in cytoplasmic quality control

**DOI:** 10.1101/2022.11.11.516101

**Authors:** Luisa Querl, Yen-Yun Lu, Christian Beißel, Heike Krebber

## Abstract

Accurate translation requires correct mRNAs with intact open reading frames. Cells eliminate defective transcripts to prevent mistranslation by three cytoplasmic mRNA quality control events termed nonsense-mediated decay (NMD), no-go decay (NGD) and non-stop decay (NSD). Translation termination on correct transcripts requires Dbp5 (human DDX19), which delivers eRF1 to the ribosomes and prevents an early contact of eRF1 with eRF3, precluding the immediate dissociation of both release factors and subsequent termination readthrough. Here, we report evidence for an influence of Dbp5 on NMD, as it delivers eRF1 also to PTC-containing transcripts. In contrast to regular translation termination and NMD, functional NGD and NSD require the eRF1-eRF3-like proteins Dom34-Hbs1. We suggest that Dbp5 delivers Dom34 to NGD and NSD substrates as well. However, in contrast to regular termination, it does not prevent an Hbs1 contact, but allows formation of a ternary Dom34-Hbs1-Dbp5 complex. The Dbp5-mediated delivery of Dom34-Hbs1 in NGD and NSD might rather shield and position the complex to prevent a premature contact of Dom34 and Rli1 to prevent inefficient splitting of the ribosomal subunits. Together, we have gathered evidence suggesting an important role of Dbp5 in cytoplasmic mRNA quality control.

## INTRODUCTION

Eukaryotic mRNAs have to complete several different quality control checkpoints to ensure the production of correct polypeptides. In nuclear mRNA quality control, the transcript is monitored for correct splicing, 5’ capping, and 3’ polyadenylation, mediated by a small number of guard proteins that retain the transcripts until processing is completed and then allow the export receptor Mex67 to bind for the export into the cytoplasm (1-6). On defective transcripts, Mex67 binding is prevented and the guard proteins recruit the degradation machinery instead.

The cytoplasmic quality control, on the other hand, controls the intactness of the transcript’s open reading frame (ORF) and therefore requires the ribosome. In case a premature stop codon (PTC) is detected, such RNAs are eliminated by nonsense-mediated decay (NMD). When the ribosome stalls on a regular codon, such transcripts are eliminated by the no-go decay (NGD) and the non-stop decay (NSD) pathways (1-4,7,8). Ribosome stalling on normal codons can originate from transcripts with rare codons or from mRNAs that contain strong secondary structures such as stem loops. These transcripts are eliminated by NGD. RNAs that lack a stop codon altogether are eliminated by NSD. Often, these are either fractured mRNAs on which ribosomes stall, or normal mRNAs lacking a stop codon entirely. On such RNAs, the ribosome translates into the poly(A) tail, which alerts the NSD machinery as well (8). In all three surveillance mechanisms, the false mRNA and the truncated protein are rapidly degraded to avoid an accumulation of defective and potentially toxic proteins (1,9).

In NMD, a PTC is detected by the Upf1-2-3 complex, which contacts eRF1 and eRF3 as well as several supporting factors, among them the guard proteins Gbp2 and Hrb1, to elicit transcript degradation (10-13). In yeast, NMD substrates are mostly degraded from the 5’ end via the exoribonuclease Xrn1 subsequent to Dcp1-Dcp2-mediated decapping, but can also be eliminated through 3’ degradation by the exosome (14). In NGD, the mRNA is first endonucleolytically cleaved, possibly by Cue2, in close proximity to the stalled ribosomes, resulting in two mRNA fragments with free 3’ and 5’ termini (15). The 3’ fragment is rapidly degraded by Xrn1 and the 5’ fragment is thereafter recognized as an NSD substrate. A complex of Dom34 (Pelota in mammals) and Hbs1, bound to GTP, recognizes the stalled ribosome and together with Rli1 promotes its splitting into the 40S and 60S subunits (16,17). Interestingly, Dom34 and Hbs1 are orthologues of the eukaryotic release factors eRF1 and eRF3, respectively (18-20). In contrast to eRF1, Dom34 lacks the NIKS motif required for stop codon recognition and the methylated GGQ motif required for peptide hydrolysis, which is why the ribosomal 60S subunit is still peptidyl-tRNA-bound after splitting (21). However, this unnatural connection is released by the ribosome quality control (RQC) complex, which mediates the rapid degradation of the incomplete nascent peptide (22-25).

Recently, it has been shown that regular translation termination might not only require eRF1 and eRF3 for stop codon recognition and polypeptide chain release, but also the DEAD-box RNA helicase Dbp5/Rat8 (26-28). It has been shown that the helicase recruits eRF1 in the cytosol and delivers it to the ribosome, thereby protecting it from premature eRF3 contact, which already waits at the ribosome for eRF1 arrival (26). Defects in *DBP5/RAT8* enable premature contact, leading to the dissociation of both proteins before proper termination, promoting translational readthrough (26). Since termination at a PTC also requires eRF1 and eRF3, it seems likely that the function of Dbp5 may also influence NMD, which has already been suggested (29). However, experimental support for this is currently missing. Furthermore, it is conceivable that Dbp5 additionally functions in NGD and NSD, as Dom34 and Hbs1 are orthologues of eRF1 and eRF3.

Here, we present evidence that Dbp5 does indeed impact the cytoplasmic mRNA quality control pathways. Mutations in its gene potentially result in stop codon readthrough and the accumulation of NMD targets in cells. Moreover, we found physical interactions between Dbp5 and Dom34-Hbs1. The formation of a ternary complex, which was shown in competition assays, is different from the interaction of Dbp5 with only eRF1 and not eRF3. In NGD and NSD the helicase seems to deliver both Dom34 and Hbs1 together to the stalled ribosome. We suggest a model in which nucleotide hydrolysis by Dbp5 and Hbs1 leads to an accommodation of Dom34 in the A-site of the stalled ribosome. Subsequently, a complex of Dbp5-ADP and Hbs1-GDP may dissociate from the ribosome, allowing Rli1 to contact Dom34. Their interaction might elicit splitting of the ribosome into its subunits and degradation of the mRNA and protein.

## MATERIALS AND METHODS

### Yeast strains and plasmids

All *Saccharomyces cerevisiae* strains, plasmids and oligonucleotides used in this study are listed in the expanded view tables S1, S2 and S3, respectively. The strain HKY1693 was generated by crossing HKY477 (*rat8Δ*) with HKY1631 (*DOM34-GFP*) and the different *rat8* mutants were generated by the exchange of pHK629 (*RAT8 URA*) with either pHK630 (*RAT8 LEU*), pHK638 (*rat8-3 LEU*), or pHK693 (*rat8-2-MYC LEU*). The strain HKY1889 was created by crossing HKY477 (*rat8Δ*) with HKY1797 (*dom34Δ*) and different *rat8* mutants were generated by plasmid exchange of pHK629 (*RAT8 URA*) with either pHK630 (*RAT8 LEU*), pHK638 (*rat8-3 LEU*), pHK654 (*rat8-7 LEU*), or pHK693 (*rat8-2-MYC LEU*). The strain HKY1845 was generated by crossing HKY492 (*upf1*Δ) with a strain that has an opposite mating type. The plasmid pHK1631 (*P*_*ADH1*_*MYC-HBS1*) was generated by digestion of pHK825 (*P*_*ADH1*_*3xMYC*) with *BamH*I and *Hind*III and insertion of the PCR product of HK2470 and HK2471 amplified from genomic DNA. For generation of the plasmid pHK1647 (*P*_*ADH1*_*DOM34-MYC*), the Dom34 open reading frame was amplified with HK3179 and HK3219 and inserted into the *Cla*I digested vector pHK825 (*P*_*ADH1*_*3xMYC*). For generation of the plasmid pHK1648 (*P*_*DOM34*_*DOM34-MYC*), the plasmid pHK825 (*P*_*ADH1*_*3xMYC*) was digested with *Cla*I and *Sac*II and the Gibson Assembly PCR product of HK3218 and HK3219 amplified from genomic DNA was inserted. The plasmid pHK1672 (*P*_*ADH1*_*DOM34-GFP*) was generated by digestion of pHK1335 (*P*_*ADH1*_*SUP45-GFP*) with *Cla*I and *Xho*I to remove the Sup45 open reading frame and insertion of the PCR product of HK3482 and HK3483 amplified from genomic DNA.

### Fluorescence microscopy

Microscopy experiments were performed as previously described (13). Logarithmic yeast cells were fixated with formaldehyde and immediately harvested, washed once with 0.1 M potassium phosphate buffer pH 6.5, once with P solution (0.1 M potassium phosphate buffer pH 6.5, 1.2 M sorbitol) and resuspended in P solution. The cells were incubated for 15 min on polylysine coated microscope slides, permeabilized with 0.5% Triton X-100 in P solution for approx. 1 min and washed once with P solution and once with Aby wash 2 (0.1 M Tris pH 9.5, 0.1 M NaCl). DNA was stained with DAPI (1 μg/ml in Aby wash 2) for 5 min and washed three times for 5 min with Aby wash 2. Microscope slides were dried and the cells mounted in 40% (v/v) glycerol, 20% (v/v) PBS (137 mM NaCl, 2.7 mM KCl, 10 mM KH_2_PO_4_, 2 mM Na_2_HPO_4_ and 1% (w/v) n-propyl gallate). Microscopy images were taken with a Leica AF6000 microscope and a LEICA DFC360FX camera with the LEICA AF 2.7.3.9 software. Z-stacks (10 images, 0.2 μm) were deconvoluted (blind, 10 iterations, with the LEICA AF 2.7.3.9 software).

### Cell lysis for western blot analysis

For Fig. 1E, log phase cells were lysed in SDS sample buffer (62.5 mM Tris pH 6.8, 2% (w/v) SDS, 10% (v/v) glycerol, 0.05% (w/v) bromophenol blue and 5% (v/v) 2-mercaptoethanol) with one pellet volume (or 200 μl for smaller cell pellets) of glass beads (0.4–0.6 mm) and heated at 95°C for 5 min. Glass beads and cell debris were removed by centrifugation at 16000x g for 5 min. The supernatant was used for western blotting.

**Figure 1:**
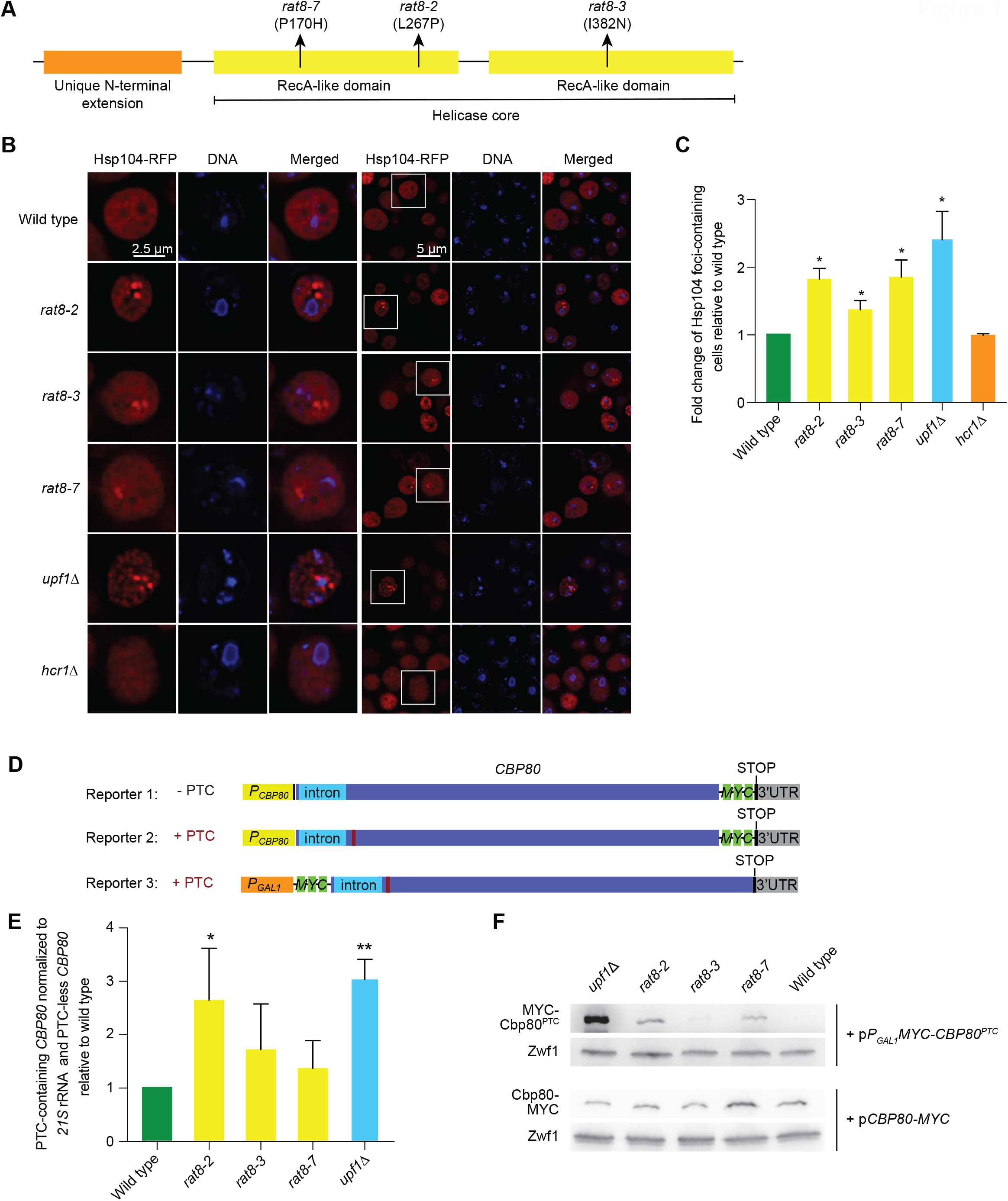
Dbp5 may be important for fully functional NMD. (A) Schematic overview of the *dbp5* mutants *rat8-2, rat8-3* and *rat8-7*. (B) Protein aggregation is increased in mutants of *RAT8*. Localization of RFP-tagged Hsp104 is shown in the indicated strains that were grown to the logarithmic growth phase at 25 °C and shifted to 37 °C or 16 °C (*rat8-7*) for 1 h. *n=3*. (C) Microscopic figures from (B) were analyzed by calculating the percentage of cells showing Hsp104 foci in each strain and relating to wild type. Results were obtained from three independent experiments and are represented as mean with standard deviation. At least 100 cells were analyzed for each strain in each experiment. *n=3*. (D) Scheme of the reporter constructs used in (E) and (F). (E) Dbp5 is required for the effective degradation of PTC-containing *CBP80* transcripts. qPCRs from total RNA of the indicated strains were carried out and reporter levels were normalized to that of *21S* rRNA. The relative abundance of PTC-containing per PTC-less reporter in wild type was set to 1 and the other strains are shown in relation. *n=4*. (E) Protein expression of the NMD reporter increased in *rat8* mutants. Expression of plasmid encoded *P*_*GAL1*_*MYC-CBP80*_*PTC*_ was monitored by western blot. Zwf1 served as a loading control. *n=2*

### Co-immunoprecipitation (co-IP) experiments

The *in vivo* co-IP studies were carried out according to the protocols published previously (29-31) For immunoprecipitation of GFP- and MYC-tagged proteins, 10 μl slurry of GFP- or MYC-Trap^®^_A beads (Chromotek) were used per reaction and incubated with 1000 μl of the cleared yeast lysate for 3 h under rotating at 8°C. Subsequently, the immunoprecipitated proteins were washed 3-5 times with PBSKMT buffer. Finally, the eluted proteins were separated on 10% SDS-polyacrylamide gels and analyzed by western blotting.

### Trichloroacetic acid (TCA) precipitation

To visualize proteins that could not be detected in lysates during co-IP experiments, precipitation with TCA was performed. Cells were harvested during log phase, resuspended in 0.1% (w/v) NaDOC and incubated at room temperature. Subsequently, 100 μl 99% (w/v) TCA were added and the samples incubated on ice for 15 min. Precipitated proteins were spun down at 13,000 rpm for 15 min and washed three times with ice-cold 80% (v/v) acetone. After air-drying the pellet, proteins were dissolved in HOECHST buffer (8 M urea, 200 mM Tris-base, 200 mM DTT, 2% (w/v) SDS, bromphenolblue), boiled at 95 °C for 5 min and loaded onto an SDS gel.

### Combined *in vivo* and *in vitro* studies

For experiments shown in Fig. 4, *in vivo* and *in vitro* protein binding systems were combined. Recombinant proteins were expressed as described earlier (26). The *in vivo* produced GFP-tagged proteins were immobilized as described in the co-IP experiments. After immobilization, the co-purified proteins were detached from the protein of interest by intense washing. For this, the proteins bound to the GFP beads were washed 5 times with PBSKMT buffer and incubated with *in vitro* lysis buffer (50 mM Tris-HCl pH7.5, 150 mM NaCl, 2 mM MgCl_2_, 5% (v/v) glycerol, 1 mM Dithiothreitol (DTT), 0.2% (v/v) NP-40 and EDTA-free protease inhibitor mix (Roche)) under agitation at 8 °C for 20 min. Subsequently, the samples were washed 3 times with *in vitro* lysis buffer. Afterwards, either purified protein or cleared bacterial cell lysate containing the expressed recombinant protein was added to the immobilized protein and incubated at 8 °C with rotation for 2 h to allow binding to the GFP-tagged protein of interest. Subsequently, the beads were washed 5 times with *in vitro* lysis buffer. Finally, the beads with the bound proteins were mixed with 30 μl of 2 x SDS buffer before the samples were loaded onto an SDS-PAGE for western blotting.

### Displacement assay

The displacement assay used in Fig. 4C is a combination of *in vivo* and *in vitro* experiments. After binding of the recombinant protein to the *in vivo* immobilized MYC-tagged protein, the beads were washed as described in the combined *in vivo* and *in vitro* studies. Subsequently, yeast lysate containing the protein of interest was added to the beads and incubated at 8 °C with rotation for 1.5 h. This protein was added in increasing concentrations to investigate a possible competition with previously formed interactions. Finally, the beads were washed 3 times with *in vitro* lysis buffer and the samples were applied to SDS-PAGE and western blotting.

### Western blot analyses

Nitrocellulose membranes were cut into pieces that compose either higher, medium or lower molecular weight proteins, allowing the simultaneous detection with different antibodies. Polyclonal rabbit antibodies were used in the following concentrations: Dbp5 (dilution 1:3000), for uS3 = Rps3 (anti-Rps3 peptide antibody 1:3000), Zwf1 (dilution 1:4000), Asc1 (dilution 1:1000). GFP-tagged proteins were detected with the anti-GFP antibody GF28R from Pierce Protein Biology (1:5000), MYC-tagged proteins with an anti-MYC antibody (sc-789; Santa Cruz, dilution 1:750). Secondary anti-rabbit IgG (H+L)-HRPO and anti-mouse IgG (H+L)-HRPO (Dianova) antibodies were used and detected with Amersham ECL Prime Western Blotting Detection Reagent (GE Healthcare) and the FUSION-SL chemiluminescence detection system (Peqlab).

### Quantitative polymerase chain reaction (qPCR)

qPCR primer pairs were designed with the GeneScript Real-time PCR (TaqMan) primer designer (Genescript.com) to reach amplification efficiencies of 0.8 -1. All samples, except the minus reverse transcriptase (-RT) and negative controls, were prepared in triplicates. Each sample was prepared in 10 μl volume consisting of 5 μl 2x qPCR master mix (qPCRBIO SyGreen Mix Lo-Rox, Nippon genetics), 4μl cDNA (0.5 ng/μl), 0.08 μl (1000 ng/μl) of forward and reverse primers and 0.86 μl nuclease free water. The qPCR was executed in a two-step protocol with 95 °C initial denaturation and 45 cycles with 30 s at 60 °C and 10 s at 95 °C. For the verification that only one specific product was amplified, the qPCR cycler CFX connect (BioRad) recorded a melting curve starting from 65 °C to 95 °C. The target RNAs were normalized to the *21S* rRNA and calculated using the 2^−ΔΔCq^ method (Livak and Schmittgen, 2001).

### Statistical analysis

The number of independent biological replicates performed for each experiment is indicated. The mean ± standard deviations are shown. All data were analyzed for significance by unpaired, two tailed, two-sample unequal variance Student’s t-test. Significant P-values below 0.05 were indicated by asterisks (*P<0.05; **P<0.01, ***P<0.001).

## RESULTS

### Dbp5 might be involved in NMD

We have shown that Dbp5 is involved in translation termination by preventing a premature eRF1-eRF3 interaction (26,28). This function implies that Dbp5 may potentially have an impact on NMD as well, as NMD-mediated termination also depends on eRF1 and eRF3 (29). A first indication for such an involvement can be investigated through the detection of protein aggregates, indicated by Hsp104 foci in the cytosol of mutants impaired in cytoplasmic quality control (32). In wild type cells, protein aggregation rarely occurs, leading to few cells exhibiting foci (32) (Fig. 1B). In a *UPF1* deletion strain, however, the number of cells displaying Hsp104 foci increases because more defective RNAs producing defective proteins accumulate (32) (Fig. 1B, C). We investigated whether the mutation of *DBP5/RAT8* would also lead to such defects and an increase of Hsp104 foci. Indeed, all mutants of *DBP5* tested, namely *rat8-2, rat8-3 and rat8-7* (33) and Fig. 1A, showed an increase of cells that contained Hsp104 foci at the non-permissive temperatures (Fig. 1B, C). In comparison, the deletion of the translation factor Hcr1 did not result in an increase of cells exhibiting protein aggregation (Fig. 1B, C). To investigate whether these aggregates present in the mutant *dbp5* strain are indeed generated by defective NMD, we carried out an NMD assay. For this we used reporter constructs that we previously created for study of NMD in yeast (13) (Fig. 1D). The NMD reporter carries the intron-containing *CBP80* gene, expressed from its own promoter as a C-terminally MYC-tagged protein (Fig. 1D, Reporter 2). It contains a PTC after the intron sequence, causing the RNA to be eliminated and its protein expression to be prevented through NMD in wild type cells (13) (Fig. 1E, F). The control reporter has no PTC but is otherwise identical (Fig. 1D, Reporter 1). We detected the amount of the PTC-containing reporter RNA via qPCR and related the results to that of the PTC-less control reporter and mitochondrial *21S* rRNA to study PTC-, NMD-specific effects. Like previously demonstrated, the PTC-containing reporter showed an increased level in the *upf1*Δ strain (Fig. 1E). We analyzed the three mutants of *DBP5/RAT8* in relation to wild type and found an increase of the PTC-containing RNA in *rat8-2* after shifting the strain to 37 °C for 1 h. Combined with the large increase of cells displaying Hsp104 foci, it appears that this mutant is affected by an impaired NMD pathway (Fig. 1E). This could suggest that when Dbp5 is substantially defective, NMD also becomes deficient. To investigate whether protein expression of the PTC-containing reporter also increases in *dbp5* mutants, which would support the main purpose of NMD to prevent aberrant protein expression, we exchanged the endogenous promoter to a galactose-inducible reporter (Fig. 1D, Reporter 3) and determined protein expression on western blots. We found that while the protein expression was low in wild type and high in *upf1*Δ cells, *rat8-2* showed an increased protein expression level as well (Fig. 1E, F). Together, these findings suggest that the proper function of Dbp5 may be important for fully effective NMD.

### Dbp5 appears to be involved in no-go and non-stop decay

Although protein aggregates can be generated through defective NMD, it is important to note that defects in other cytoplasmic quality control pathways can contribute to protein aggregation as well. Mutation of the non-stop and no-go decay factor Dom34, for instance, also results in the formation of Hsp104 foci (32) (Fig. 2A). Given the similarities between Dom34-Hbs1 and eRF1-eRF3, it is conceivable that Dbp5 might not only deliver eRF1 to regular stop codons, but also carry Dom34 to stalled ribosomes that are arrested on normal codons, which become substrates for NGD and NSD. To investigate this, we first carried out the Hsp104-RFP microscopy experiment and analyzed protein aggregation in the double mutants of *dom34*Δ *rat8*. We had observed that each *rat8* mutant alone as well as *dom34*Δ led to increased Hsp104 foci (Fig. 1B, C, 2A, B). Interestingly, the effects did not fully add up when the mutations were combined and all double mutants showed a similar or only slightly higher degree of protein aggregation in comparison to the single mutants (Fig. 1B, C, 2A, B). This implies that effects from single *rat8* mutants overlap in part with that of *dom34*Δ and supports the possibility that Dbp5 may act in the same pathway as Dom34. To further investigate this, we examined a possible connection between *DBP5* and *DOM34* or *HBS1* in a *rat8-2* strain transformed with plasmids overexpressing the NSD- and NGD factors by growth analysis at different temperatures, but could not verify a reproducible genetic interaction (Fig. 2C).

**Figure 2:**
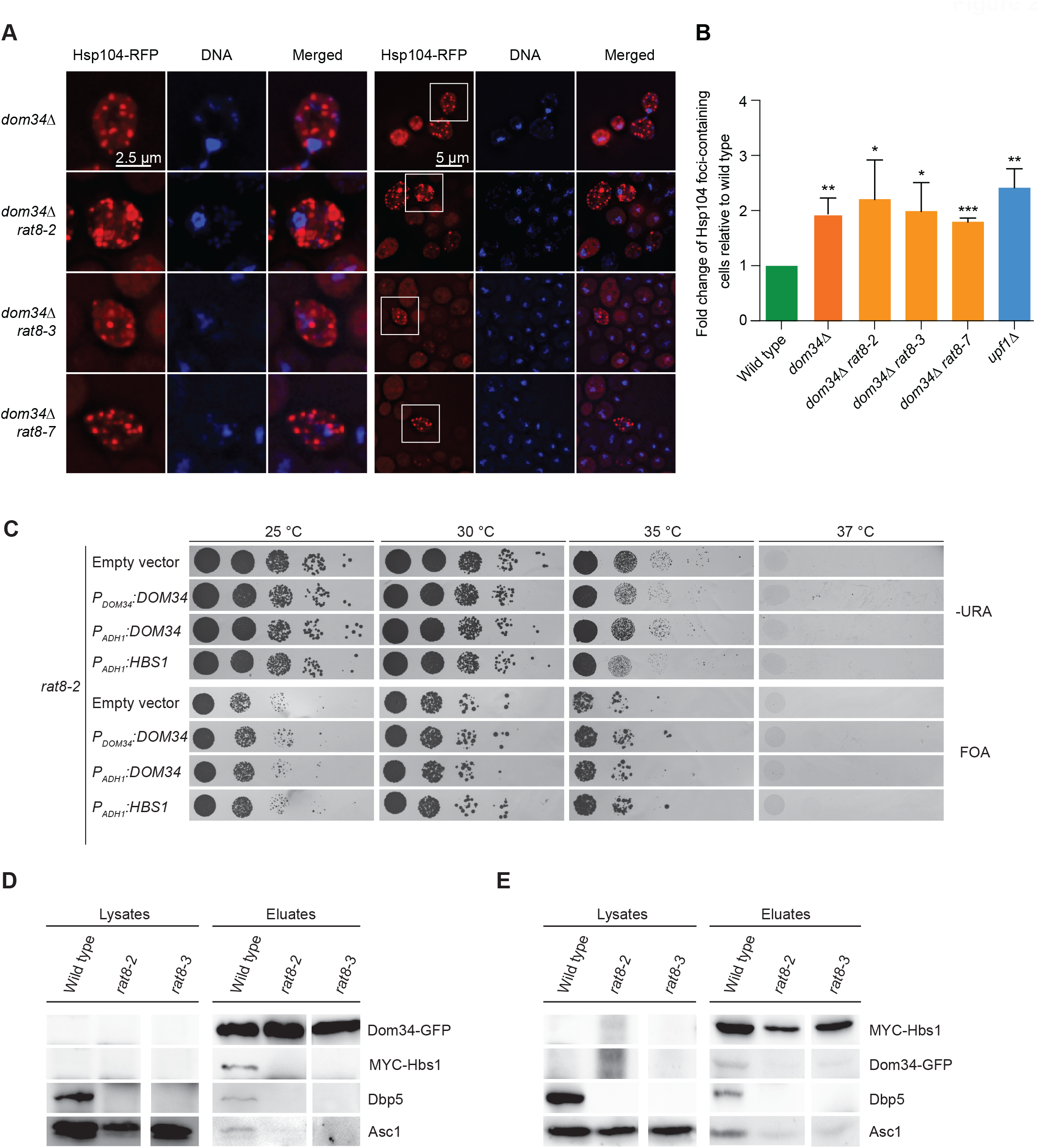
Dbp5 participates in NGD and NSD. (A) Protein aggregation is not increased in the double mutants of *dom34Δ rat8* as compared to the single mutants. Localization of RFP-tagged Hsp104 is shown in the indicated strains that were grown to the logarithmic growth phase at 25 °C and shifted to 37 °C or 16°C (*rat8-7*) for 1 h. *n=4*. (B) Microscopic figures from (A) were analyzed as in Fig. 1B. Cells that contained Hsp104-RFP foci were counted and the fold change increase of cells with aggregates is shown. All results were normalized to the *DBP5/RAT8* wild typical strain and the error bars represent the standard deviations between different experiments. *n=4*. (C) Growth analysis of 10-fold serial dilutions of the indicated strains transformed with the indicated plasmids (containing the *URA3* selection gene) is shown. Growth without the plasmids is shown on FOA plates. All plates were incubated for 3 days at the indicated temperatures. *n=3*. (D) The interaction of Dom34 with the ribosome and with Hbs1 depends on functional Dbp5. Western blot analysis of Dom34 co-IPs in wild type, *rat8-2* and *rat8-3* cells shifted to 37 °C for 1 h is shown. All strains contained plasmid encoded *MYC-Hbs1*. The co-precipitated Asc1 protein indicates ribosome binding. *n=3*. (E) The interaction of Hbs1 with the ribosome and Dom34 is decreased in *rat8* mutants. Western blot analysis of the co-precipitation of Dom34, Asc1 and Dbp5 with Hbs1 in different strain backgrounds is shown. *n=3*.

Since *DBP5* and *DOM34* or *HBS1* appeared to not interact genetically, we focused on monitoring possible physical interactions between the proteins. During translation termination, Dbp5 can be found in a complex with eRF1 that does not contain eRF3 (26). Since eRF1 and eRF3 are structurally very similar to their orthologues Dom34 and Hbs1 (18-20), we analyzed their binding to Dbp5 utilizing co-IP experiments. GFP-tagged Dom34 was precipitated from yeast lysates and the co-purification of MYC-tagged Hbs1 and Dbp5 is shown on western blots (Fig. 2D). Interestingly, while both proteins interact with Dom34, their association is disrupted in mutant *rat8-2* and *rat8-3* cells (Fig. 2D). The same result was obtained when we precipitated Hbs1. Dom34 and Dbp5 were co-precipitated with Hbs1 in wild type cells, but not in the *rat8* mutants (Fig. 2E), suggesting formation of a Dom34-Hbs1-Dbp5 complex that requires functional Dbp5. Further examination of the ribosomal protein Asc1 revealed its co-purification with both Dom34 and Hbs1 in wild type cells, which was lost in the *rat8* mutants (Fig. 2D, E), indicating that functional Dbp5 is also important for the association of the Dom34-Hbs1 complex with the ribosome.

Together, these results suggest that Dbp5 is involved in the same pathway as Dom34 and Hbs1. It appears that Dbp5 is able to form a physical complex with both proteins, however, in contrast to normal translation termination, in which eRF3 is absent from the Dbp5-eRF1 complex, the Dom34-Dbp5 complex seems to also contain Hbs1. Furthermore, we show that the interaction of Dom34 with Hbs1 and their association with the ribosome depends on functional Dbp5.

### Dom34, Hbs1 and the ATP-bound form of Dbp5 physically interact

Dbp5 recycling occurs at the NPC, where it requires Nup159/Rat7 to be able to newly bind ATP (27,34,35). To investigate whether recycling of Dbp5 is required for the formation of the ternary Dbp5-Dom34-Hbs1 protein complex, we used the mutant *rat7ΔN*, which lacks the N-terminus that mediates contact with Dbp5 (36). In this mutant background, the Dbp5 protein is intact but not efficiently recycled to ATP-bound Dbp5 anymore. Clearly, while under wild typical conditions Hbs1, Dbp5 and Asc1 co-precipitated with Dom34, these interactions were abolished in the absence of recycled Dbp5 (Fig. 3A). To show that the proteins of interest were indeed expressed despite weak lysate signals, we performed trichloroacetic acid precipitation to increase protein concentration and could show that the proteins were indeed expressed in *rat7*ΔN (Fig. 3B).

**Figure 3:**
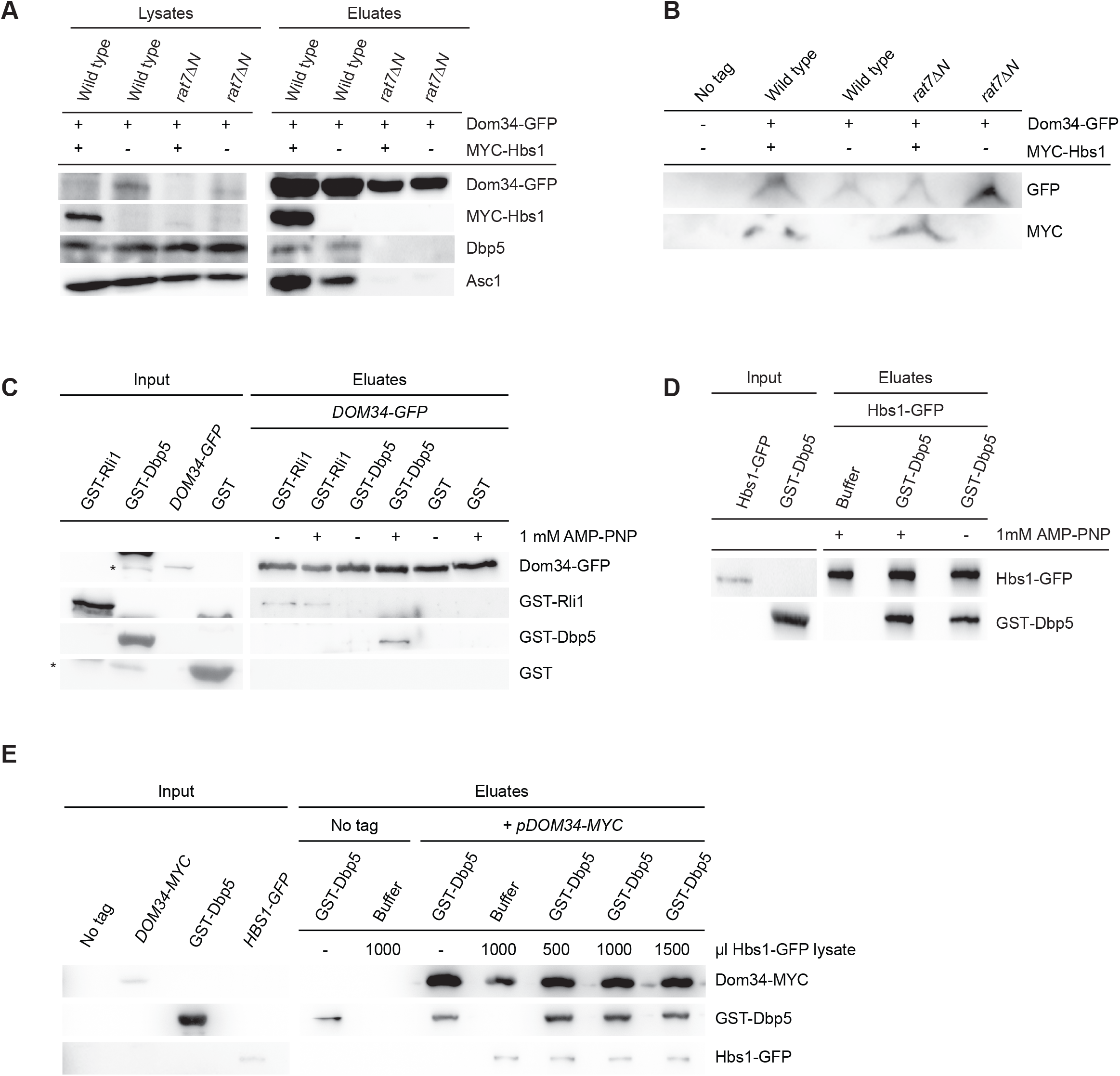
Functional Dbp5 ATPase activity is required for the Dom34-Hbs1 interaction. (A) Complex formation of Dom34 with Hbs1, Dbp5 and the ribosome depends on recycled, ATP-bound Dbp5. Western blot analysis of the co-IP experiment of Dom34 shows a reduced co-precipitation of Hbs1, Dbp5 and Asc1 in *rat7ΔN* cells compared to wild type. *n=3*. (B) Trichloroacetic acid (TCA) precipitation was performed for lysate samples of (A) and the protein signals are shown on a western blot. (C) Dbp5 and Dom34 directly interact in the presence of a non-hydrolysable ATP analogue. A combined *in vivo* and *in vitro* binding study was carried out with Dom34-GFP purified in pull-down experiments from yeast cell lysates to which recombinant GST-Dbp5 or GST-Rli1 and, if indicated, 1 mM AMPPNP were added. Western blot analysis reveals direct protein interactions. Asterisks indicate bands that likely arose from slight levels of protein degradation. *n=3*. (D) The direct protein interaction of Dbp5 and Hbs1 is independent of the nucleotide status of Dbp5. Endogenously produced Hbs1-GFP was pulled down with GFP-Trap® beads and incubated with recombinant GST-Dbp5 supplemented with or without AMPPNP. Direct protein interaction was confirmed by western blot analysis. *n=3*. (E) A preformed complex of Dom34 and Dbp5 was able to interact with Hbs1. Western blot analysis of a competition assay of the indicated endogenously and recombinantly expressed proteins is shown. Increasing amounts of Hbs1 were added to a preformed complex of GST-Dbp5 and Dom34-GFP. 1 mM AMPPNP was supplemented to the buffer during the GST-Dbp5 incubation step. *n=3*.

To further investigate the binding requirements for the new Dbp5 interaction partners, we carried out combined *in vivo* and *in vitro* studies. We first immobilized Dom34-GFP from whole yeast lysates (*in vivo*) to beads and subsequently added recombinant GST-Dbp5 for *in vitro* binding. To analyze whether ATP is required for the interactions, we performed the experiment with or without 1mM AMPPNP. As shown in Fig. 3C, Dom34 and Dbp5 showed a clear interaction in the presence of the ATP analogue, while this interaction was not detectable in its absence. In contrast to that, we show that Rli1 and Dom34 interact with each other independently of the nucleotide-bound state of Rli1 (Fig. 3C).

In an additional experiment, we investigated the nature of Hbs1 binding to Dbp5. This time we immobilized Hbs1-GFP from whole yeast lysates to beads and added recombinant GST-Dbp5. Like for Dom34, we were able to show a direct interaction of the two proteins. However, contrary to the Dom34 interaction, the Hbs1 contact with Dbp5 does not depend on ATP-bound Dbp5 (Fig. 3D). It seems likely that Dbp5 forms a complex with Dom34-Hbs1 in the ATP-bound state, but upon ATP hydrolysis dissociates from Dom34 together with Hbs1.

To find out whether the binding of Dbp5 and Hbs1 to Dom34 is mutually exclusive, we immobilized Dom34-GFP from yeast and added GST-Dbp5 in the presence of AMPPNP to generate a preformed complex. Afterwards we added increasing concentrations of yeast lysate containing MYC-tagged Hbs1. The positive controls showed that Dbp5 and Hbs1 alone were able to bind to Dom34, respectively (Fig. 4C). Interestingly, even high concentrations of Hbs1 did not disrupt the preformed complex of Dbp5 and Dom34. These experiments rather show that the proteins form a trimeric complex (Fig. 4C). These findings contrast the Dbp5-eRF1 complex formed during regular translation termination where eRF3 is missing (26).

**Figure 4:**
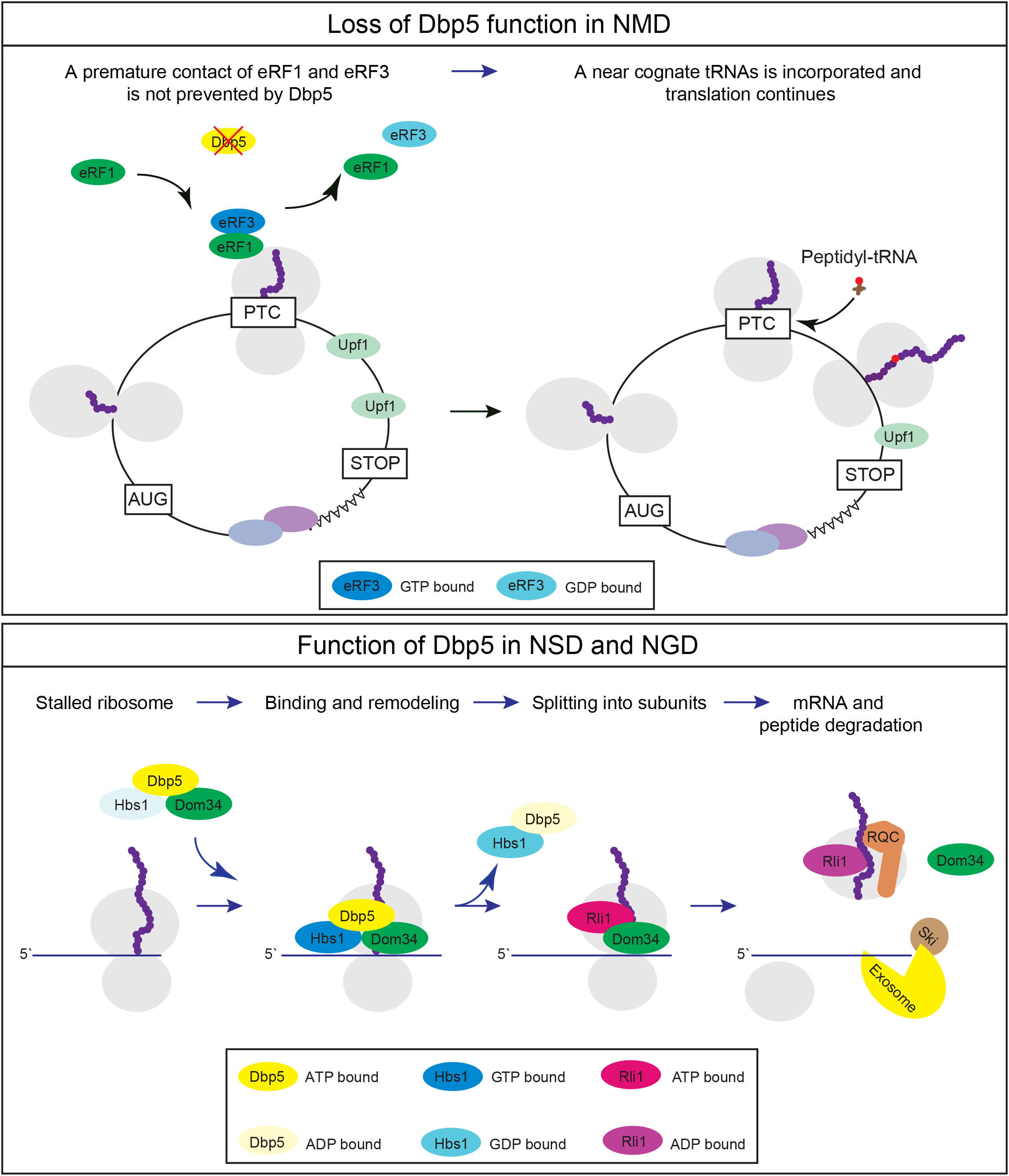
Model for the role of Dbp5 in NMD, NSD and NGD. Top: (adapted from (29)) PTC-containing mRNAs are usually terminated at the PTC prematurely via the Dbp5-mediated delivery of eRF1 to eRF3. This premature termination event is subsequently recognized by Upf1, which initiates NMD-mediated degradation of the transcript. However, mutations in DBP5 or its downregulation may lead to the premature contact of eRF1 and eRF3 at the ribosome at such PTCs, which instantly leads to the GTP hydrolysis of eRF3, resulting in the dissociation of eRF1, eRF3 and the ribosome. This enables near cognate tRNAs to enter the ribosome, resulting in the readthrough of the PTC, continued translation and suppression of NMD. Bottom: Ribosome stalling occurs on broken mRNAs, mRNAs with rare codons or stem loop structures and on mRNAs that lack a stop codon. These ribosomes cannot be removed by eRF1 and eRF3, but require Dom34-Hbs1, which recognize Hel2-mediated ubiquitinylation of the stalled ribosome. Dom34 and Hbs1-GTP are delivered to the A-site of the ribosome by the helicase Dbp5-ATP. Presumably, nucleotide hydrolysis of Dbp5 and Hbs1 accommodates Dom34 in the active site and leads to the subsequent dissociation of Dbp5-ADP and Hbs1-GDP, allowing Rli1 to bind to Dom34. This contact triggers the ATP hydrolysis of Rli1 to split the ribosome into its subunits. The mRNA is rapidly degraded by the Ski complex and the exosome and the non-released truncated protein is degraded via the ribosome quality control (RQC) pathway.

Based on our results, we suggest that Dbp5 is involved in the cytoplasmic mRNA quality control systems. It may promote proper activation of NMD through its role in eRF1-eRF3-mediated termination and acts in NSD and NGD by facilitating Dom34-Hbs1 recruitment (Fig. 4). For the latter process, we propose a model in which Dom34, Hbs1-GTP and Dbp5-ATP form a complex that enters stalled ribosomes. Subsequent ATP- and GTP-hydrolysis likely induces the remodeling of the complex, allowing Dbp5 and Hbs1 to dissociate and Dom34 to interact with Rli1 to assist splitting of the ribosome into the ribosomal subunits.

## DISCUSSION

The cytoplasmic mRNA quality control systems rely on the ribosome that is required for the recognition of defective open reading frames. Since the helicase Dbp5 has been shown to deliver eRF1 to terminating ribosomes (26,28), it may be conceivable that this mechanism operates similarly in NMD, when PTCs stop the ribosomes prematurely (29). Here, we have shown that mutations in *DBP5/RAT8* lead to an increased production of defective proteins, as increased amounts of Hsp104-RFP foci were detected (Fig. 1A, B). In addition to that, we have shown that a PTC-containing reporter transcript accumulates in the *rat8-2* mutant and translation of this transcript increases (Fig. 1D, E), which suggests that functional Dbp5 may be required for fully functional NMD. For the other *dbp5* mutants *rat8-3* and *rat8-7* no significant increase in the accumulation of the reporter construct could be detected. This is consistent with previous findings by Gross et al. (2007), which have concluded that mutations in *dbp5* do not stabilize NMD substrates (28). It appears that dysfunctional Dbp5 may have a minor adverse effect on NMD, which is further supported by the observation of increased readthrough of the termination codon in *rat8-2* (26), since readthrough has been shown to counteract NMD (37,38). This way, the loss of functional Dbp5 could result in a premature contact of eRF1 and eRF3, resulting in inefficient termination at a PTC and therefore readthrough and inefficient NMD (Fig. 4).

The other two known cytoplasmic mRNA quality control systems are NSD and NGD. In both systems, the ribosome stalls on regular codons and needs to be released from the mRNAs, which involves the two eRF1-eRF3 like proteins, Dom34-Hbs1 (9,21). Since Dbp5 appears to mediate the interaction between eRF1 and eRF3 (26), it is conceivable that the helicase also interacts with their structural counterparts. We have shown that Dbp5 and Dom34 function in the same pathway, since protein aggregation due to their mutation or deletion is not additive and a direct physical interaction between the proteins has been observed (Fig. 2 A, B, D, E). Importantly, the binding of Dom34 to Dbp5, Hbs1 and the ribosome was abolished in the *DBP5* mutants *rat8-2* and *rat8-3* (Fig. 2D). Vice versa, Hbs1 pull-down experiments showed the same loss of interactions between these proteins in *DBP5* mutants (Fig. 2E).

In addition to rescuing stalled ribosomes during NGD and NSD, Dom34-Hbs1 are also involved in the splitting of inactive stored 80S-like ribosomes, required for restarting translation after stress (39). Interestingly, van den Elzen et al. have discovered that a mutation in the binding domains of each protein to the other results in different phenotypes and they therefore speculated that an additional factor that helps to separate 80S-like ribosomes might be missing (39). Our data suggest that this missing factor might be Dbp5.

The function of the DEAD-box RNA helicase Dbp5 in regular translation is to deliver eRF1 to the terminating ribosome and by this protect it from interaction with eRF3 to prevent a premature contact, which leads to premature dissociation of the proteins and ultimately in readthrough of the termination codon (26). Interestingly, this mode of action was not detected for Dom34 and Hbs1. It appears that Dbp5 is able to simultaneously bind both proteins (Fig. 3E) and we suggest that it delivers both factors to the ribosome at the same time (Fig. 4). This seems possible because Hbs1 requires only its second domain for its interaction with Dom34, while eRF3 requires domain 2 and 3 for the eRF1 contact (39). Furthermore, Dbp5 recycling is required for these interactions (Fig. 3A), suggesting that ATP-bound Dbp5 forms a complex with Dom34-Hbs1 that interacts with the ribosome. Interestingly, immobilized Hbs1 binds to recombinant Dbp5 independently of the nucleotide status of Dbp5 (Fig. 3D), while the binding of immobilized Dom34 was only detectable if 1mM AMPPNP was present (Fig. 3C). These findings suggest that ATP hydrolysis leads to the dissociation of Dbp5-Hbs1 from Dom34, leaving Dom34 bound to the ribosome. This would suggest that in NGD and NSD, Dbp5 may not protect Dom34 from a premature contact with Hbs1, but rather with Rli1. In case of a premature contact, the proteins might not be placed properly for efficient ribosome splitting. Preventing an early contact between Dom34 and Hbs1 might either not be necessary or, due to the different binding mode, Dbp5 itself can keep the proteins apart until the stalled ribosomes are reached. Upon arrival at the ribosome, Dom34 and Hbs1 dissociate and pave the way for the Dom34-Rli1 interaction, necessary for ribosome splitting. Such a model is supported by the finding that the binding of Rli1 to Dom34 was independent of its ATP bound state (Fig. 3C). We suggest that the dissociation of Dbp5-Hbs1 enables Dom34 to bind to Rli1 for ribosome splitting, after proper positioning.

Together, our results suggest that the newly discovered function of Dbp5 in NSD and NGD is somewhat similar to that in regular translation termination. In both cases, the helicase delivers the proteins eRF1 and Dom34 to the ribosomes. However, in normal termination events, Dbp5 prevents an early eRF1-eRF3 contact, while in NSD and NGD a ternary complex of Dbp5, Dom34 and Hbs1 is formed (Fig. 2 D, E, Fig. 3E). Together with our biochemical data, we suggest a model in which Dbp5 delivers Dom34 and Hbs1 to stalled ribosomes. ATP hydrolysis of the helicase Dbp5 and GTP hydrolysis of Hbs1 result in their dissociation from the ribosome, allowing Dom34 to contact Rli1 for ribosome splitting (Fig. 4). Thus, Dbp5 functions not only in regular translation, but also in the cytoplasmic mRNA quality control events.

## ACKNOWLEDGEMENTS

We thank Lena Söldner and Christina Müller for technical assistance. We are grateful to T. Inada, R. Lill, U. Mühlenhoff and O. Valerius for providing antibodies, strains and/or plasmids.

## SUPPLEMENTARY INFORMATION

Table S1

*Saccharomyces cerevisiae* strains used in this study, related to Figures 1-3.

Table S2

Plasmids used in this study, related to Figures 1-3.

Table S3

Oligonucleotides used in this study, related to the methods section.

**Table S1.**
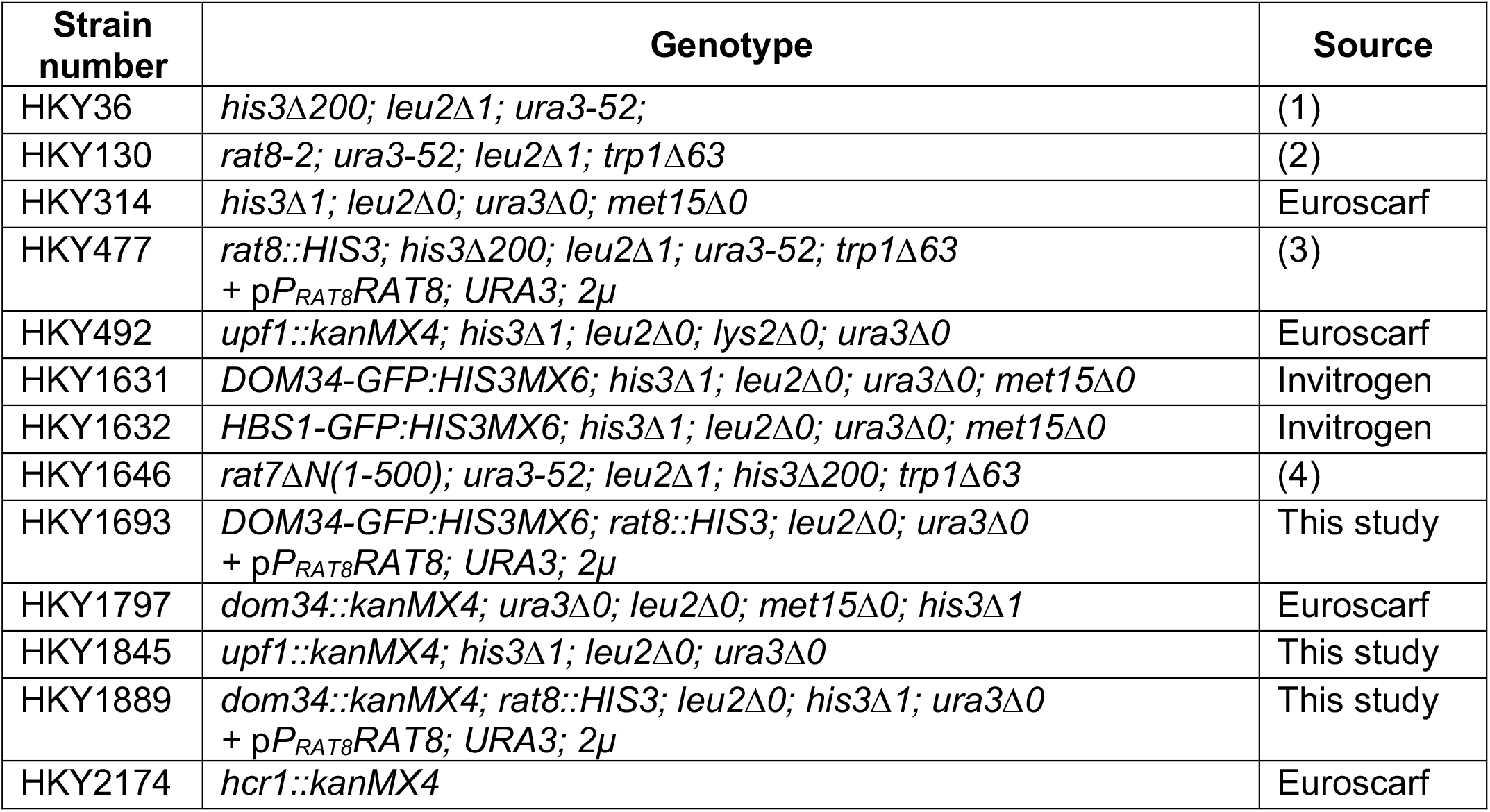
*Saccharomyces cerevisiae* strains used in this study.

---

**Table S2.** Plasmids used in this study.

| Plasmid number | Features | Source |
| --- | --- | --- |
| pHK88 | *URA3; CEN; AMP<sup>R</sup>* | (1) |
| pHK630 | *P<sub>RAT8</sub>RAT8; LEU2; 2μ; AMP<sup>R</sup>* | (2) |
| pHK638 | *P<sub>RAT8</sub> rat8-3; LEU2; CEN; AMP<sup>R</sup>* | (2) |
| pHK654 | *P<sub>RAT8</sub> rat8-7; LEU2; CEN; AMP<sup>R</sup>* | (2) |
| pHK693 | *P<sub>RAT8</sub> rat8-2-MYC; LEU2; CEN; AMP<sup>R</sup>* | (2) |
| pHK1282 | *P<sub>TAC</sub>GST-DBP5 in pGEX-4T-1; KAN<sup>R</sup>* | (3) |
| pHK1288 | *P<sub>TAC</sub>GST in pGEX-6P-1; AMP<sup>R</sup>* | (3) |
| pHK1414 | *P<sub>TAC</sub>GST-RLI1 in pGEX-6P-1; AMP<sup>R</sup>* | (3) |
| pHK1574 | *P<sub>CBP80</sub>CBP80-MYC; URA3; CEN; AMP<sup>R</sup>* | (4) |
| pHK1578 | *P<sub>CBP80</sub>CBP80<sup>PTC</sup>-MYC; URA3; CEN; AMP<sup>R</sup>* | (5) |
| pHK1600 | *P<sub>GAL1</sub>MYC-CBP80<sup>PTC</sup>; URA3; CEN; AMP<sup>R</sup>* | (5) |
| pHK1631 | *P<sub>ADH1</sub>MYC-HBS1; URA3; CEN; AMP<sup>R</sup>* | This study |
| pHK1643 | *HSP104-mRFP; URA3; CEN; AMP<sup>R</sup>* | (6) |
| pHK1647 | *P<sub>ADH1</sub>DOM34-MYC; URA3; CEN; AMP<sup>R</sup>* | This study |
| pHK1648 | *P<sub>DOM34</sub>DOM34-MYC; URA3; CEN; AMP<sup>R</sup>* | This study |
| pHK1672 | *P<sub>ADH1</sub>DOM34-GFP; URA3; CEN; AMP<sup>R</sup>* | This study |

**Table S3.**
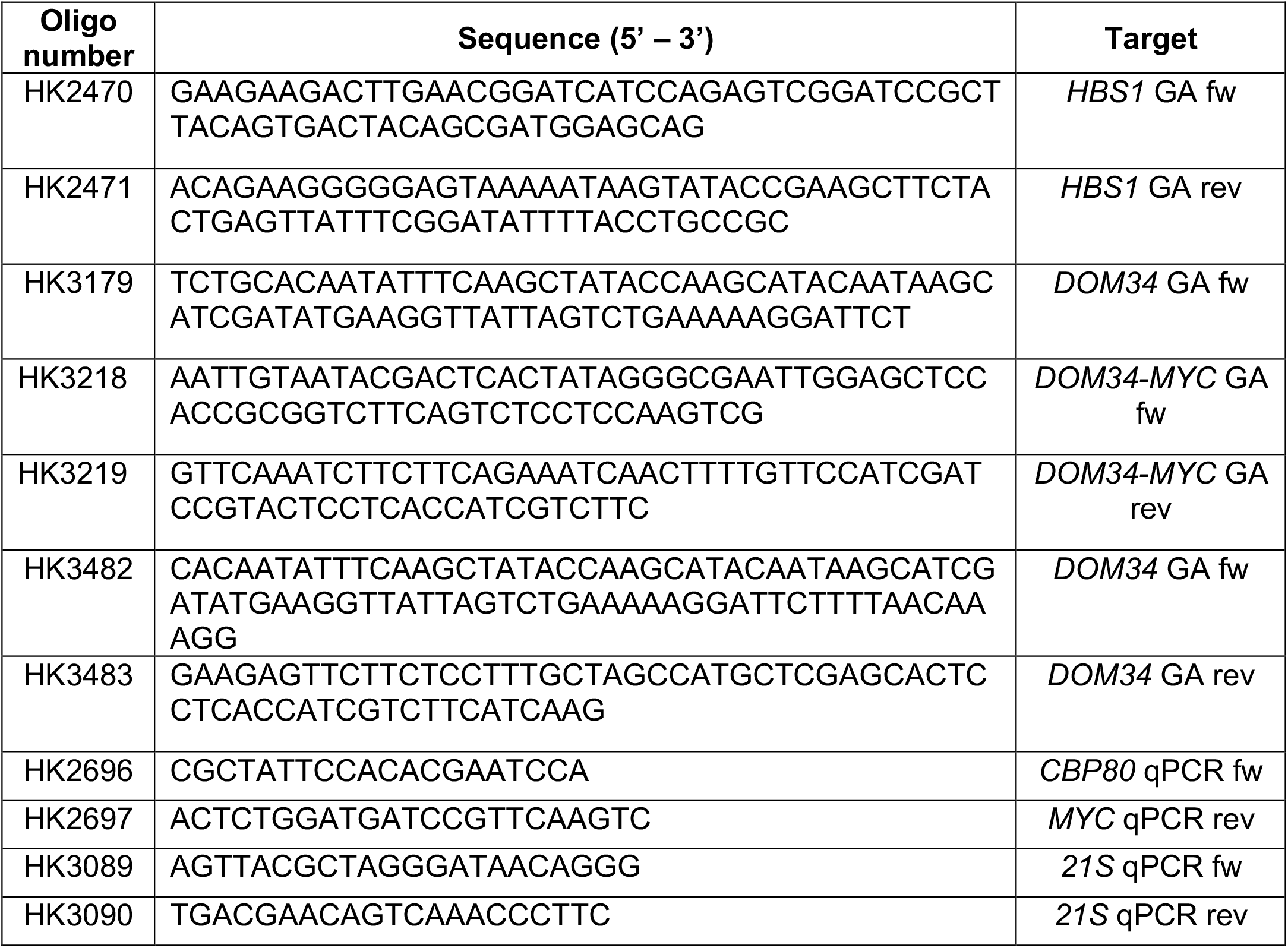
Oligonucleotides used in this study.

